# Growth-coupled continuous directed evolution by MutaT7 enables efficient and automated enzyme engineering

**DOI:** 10.1101/2024.12.16.628732

**Authors:** Yijie Deng, Kai Etheridge, Xinping Ran, Hannah E. Maurais, Rahul Sarpeshkar

## Abstract

Traditional directed evolution is limited by labor-intensive iterative steps and low-throughput selection and screening. To address these challenges, we developed a growth-coupled continuous directed evolution (GCCDE) approach, enabling automated and efficient enzyme engineering. By linking enzyme activity to bacterial growth and utilizing the MutaT7 system, GCCDE combines *in vivo* mutagenesis and high-throughput selection of superior enzyme variants in a single process. To validate this approach, we evolved the thermostable enzyme CelB to enhance β-galactosidase activity at lower temperatures while maintaining thermal stability. CelB activity was coupled to the growth of *E. coli*, allowing variants with improved activity to metabolize lactose more efficiently and promote faster bacterial growth in a minimal medium. Using a continuous culture system, we achieved automated mutagenesis and real-time selection of over 10^9^ variants per culture. Integrating *in vitro* and *in vivo* mutagenesis further increased genetic diversity, yielding CelB variants with significantly enhanced low-temperature activity compared to the wild type while preserving thermostability. DNA sequencing identified key mutations likely responsible for improved substrate binding and catalytic turnover. This GCCDE approach is broadly applicable for optimizing diverse enzymes, demonstrating the potential of automated continuous evolution for industrial and research applications.

**IMPORTANCE:** Enzyme engineering aims to develop enzymes with improved or novel traits, but traditional methods are slow and require repetitive manual steps. This study presents a faster, automated protein engineering approach. We utilized an *in vivo* mutagenesis technique, MutaT7 tools, to induce mutations in living bacteria and established a direct link between enzyme activity and bacterial growth. A continuous culture setup was used to enable growth-coupled high-throughput selection of better-performing variants. Bacteria with improved enzymes grew faster, selecting superior variants without manual intervention. Using this method, we engineered CelB with better performance at lower temperatures while maintaining high-temperature stability. The approach is adaptable to many enzymes. It offers a faster and more efficient solution for enzyme engineering. This system enables high-throughput mutagenesis and selection simultaneously, showing the power of automated continuous evolution to advance enzyme engineering.

## INTRODUCTION

### Introduction

Directed evolution mimics natural evolutionary processes in the laboratory and has become a powerful tool for engineering proteins and regulatory elements, with applications spanning therapeutics, diagnostics, and metabolic engineering (1, 2). By introducing genetic diversity and selecting advantageous phenotypes, this technique rapidly enhances enzyme and protein performance. However, traditional directed evolution relies on iterative rounds of *in vitro* mutagenesis, transformation, and selection or screening, which are labor-intensive, time-consuming, and limit efficiency and scalability in protein engineering (2–4).

To overcome these limitations, *in vivo* directed evolution methods have emerged, enabling continuous evolution of target proteins within living systems (3–5). Techniques such as phage-assisted continuous evolution (PACE) (6), orthogonal DNA Replication (OrthoRep) (7), and MutaT7 (8) have significantly advanced the field of protein engineering. Among these, the MutaT7 system uses a hypermutator chimera protein comprising T7 RNA polymerase fused to a cytidine deaminase (8). This system introduces C-to-T or G-to-A mutations in regions downstream of the T7 promoter. Given its simplicity and effectiveness, enhanced variants of MutaT7 have been developed to induce all possible transition mutations, further expanding its utility (9–11). Compared to in *vitro* methods, MutaT7 bypasses iterative steps like error-prone PCR and plasmid transformation, reducing time and labor and greatly accelerating the process of protein engineering (8–10).

Although *in vivo* mutagenesis has promoted the efficiency of directed evolution, high-throughput selection remains a major bottleneck. Growth-coupled directed evolution addresses this challenge by linking enzymatic activity to bacterial growth under selective pressure. Variants with enhanced activity promote faster growth, leading to their enrichment, while less functional variants are outcompeted in the cell population. Growth coupling can be achieved through several strategies, including enabling the targeted protein to complement auxotrophic deficiencies, to detoxify harmful compounds, or to regulate reporter gene expression linked to cell growth (12). This strategy enables the high-throughput selection of enzyme variants *in vivo* and facilitates automation (5, 12).

In this study, we developed a growth-coupled *in vivo* continuous directed evolution (GCCDE) system that combines MutaT7 mutagenesis with continuous culture for efficient enzyme evolution. Our approach enables rapid evolution of large variant libraries while precisely tuning selective pressures over extended periods. To demonstrate its effectiveness, we applied GCCDE to engineer the thermostable enzyme CelB from *Pyrococcus furiosus* (13, 14), a tetrameric protein with both β-glucosidase and β-galactosidase activities at high temperatures (15). Our goal was to enhance CelB activity at lower temperatures while retaining its thermostability. Through automated mutagenesis, selection, and enrichment in a continuous culture system, GCCDE successfully evolved CelB variants with designed properties. Our method is adaptable to other enzymes and highlights the power of growth-coupled *in vivo* directed evolution.

## RESULTS AND DISCUSSION

### Experimental design

A critical step in growth-coupled in vivo directed evolution is establishing a direct link between bacterial growth and the activity of a targeted enzyme. This requires a host bacterial strain lacking the native activity of the target enzyme, ensuring that bacterial growth in a defined medium depends entirely on the activity of the enzyme introduced *via* a plasmid. The growth medium must include the enzyme’s substrate, which is converted into essential nutrients for bacterial proliferation. Variants with higher enzymatic activity utilize the substrate more efficiently, promoting faster growth and enrichment of superior variants over time (Figure 1A).

**Figure 1.**
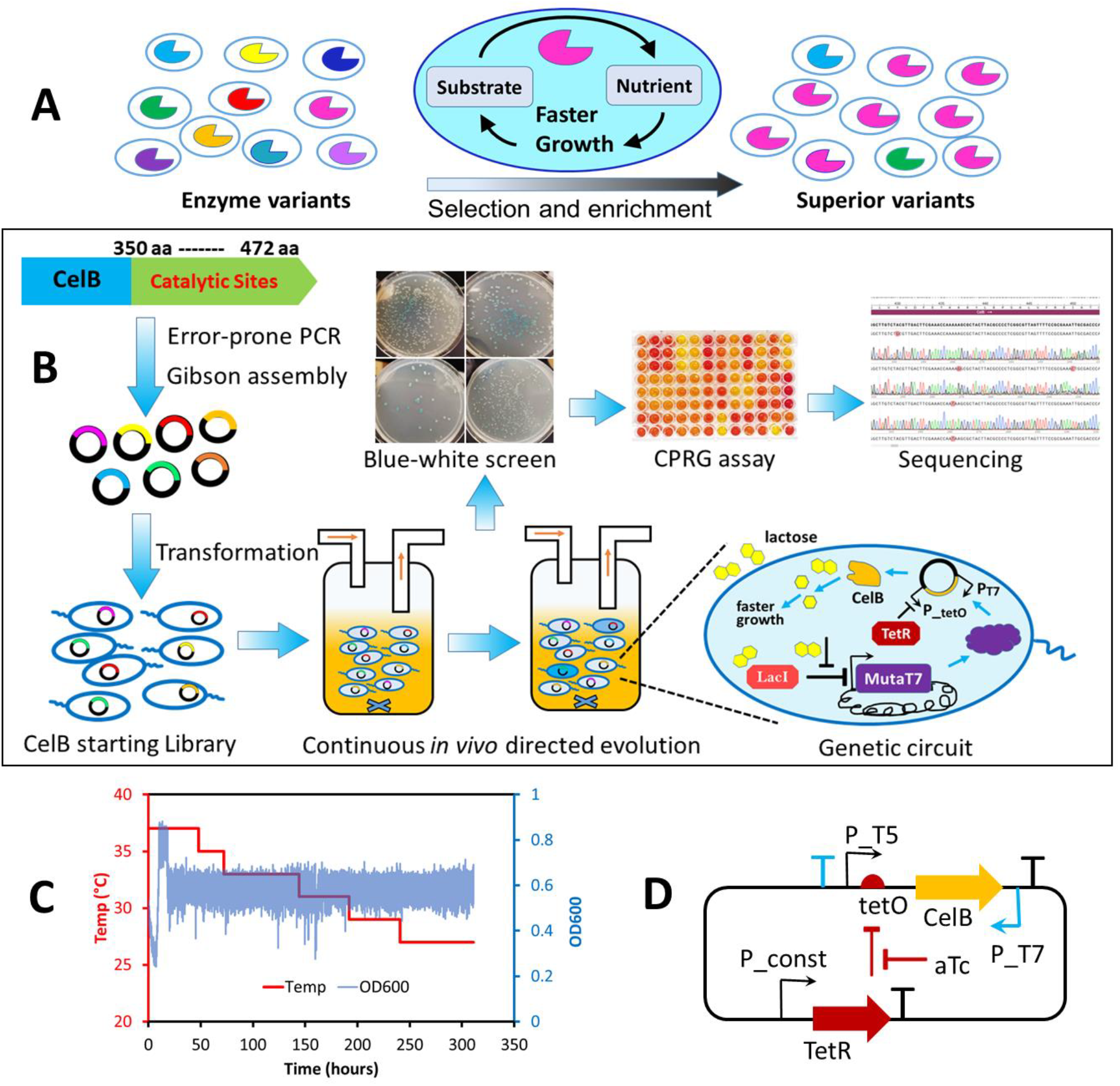
Experimental design of the growth-coupled *in vivo* continuous directed evolution. (A) Growth-coupling strategy to link enzymatic activity to cellular growth. (B) CelB starting library was generated via error-prone PCR (*in vitro* mutagenesis), followed by *in vivo* mutagenesis, and high-throughput selection in a continuous culture system. Variants with enhanced activity were isolated from blue-white screening and confirmed by CPRG assays prior to DNA sequencing. A simple genetic circuit allows for flexible control of gene expression. (C) The culture temperature was gradually reduced from 37^°^C to 27^°^C to select CelB mutants with higher activity at lower temperatures. The cell density was maintained at an OD_600_ of approximately 0.6 during the exponential phase in a turbidostat device. (D) The low-copy-number plasmid designed for this study, carrying CelB, served as both the template for error-prone PCR and the vehicle for *in vivo* mutagenesis.

As a demonstration, we targeted the thermostable CelB protein (13,14) to improve its β-galactosidase activity at lower temperatures while maintaining its stability at high temperatures. CelB activity was coupled to bacterial growth by using *E. coli* Dual7 strain (10) as the host. This strain, derived from DH10B, contains mutations in the *lacZ* gene, rendering its β-galactosidase activity negligible. The Dual7 strain also integrates MutaT7 proteins into its chromosome, along with a Δ*ung* mutation to enhance *in vivo* mutagenesis efficiency by preventing the repair of induced mutations (8,10). The strain, transformed with a plasmid library of CelB, was cultivated in a lactose minimal medium where lactose served as the sole carbon source. Enhanced CelB variants converted lactose into glucose and galactose more efficiently, enabling faster bacterial replication. Cells with less effective variants grew slower and were gradually washed out in the continuous culture system, leading to the enrichment of highly active CelB variants (Figure 1B). Selective pressure was applied by gradually lowering the culture temperature from 37^°^C to 27^°^C, favoring the evolution of CelB variants with improved activity at lower temperatures (Figure 1C).

We designed a simple genetic circuit to allow for flexible gene regulation. A low-copy-number plasmid carrying the *celB* gene under the control of the hybrid promoter P-T5-tetO allowed CelB expression to be regulated by the external inducer anhydrotetracycline (aTc) (Figure 1D). Lactose in the medium activated the expression of MutaT7 proteins, initiating *in vivo* mutagenesis. A T7 promoter downstream of the *celB* gene ensured independent regulation of mutagenesis and CelB expression, while a T7 terminator upstream of the hybrid promoter P-T5-tetO minimized off-target mutations (Figure 1D). Interestingly, the system inherently includes a beneficial negative feedback loop. As CelB activity increases and more lactose is metabolized, the reduced availability of lactose for MutaT7 expression naturally slows down mutation rates. This mechanism could help stabilize the system when the targeted protein approaches its optimal function. Although this feedback loop was not a significant factor in our current experiment (due to the high lactose concertation used in the medium saturating the LacI protein), it holds potential for enhancing future circuit designs in *in vivo* directed evolution.

To address the limitations of MutaT7, which introduces only transition mutations, we used error-prone PCR to generate a diverse starting library from the above CelB plasmid (Figure 1D) before initiating in vivo mutagenesis. The combination of *in vitro* and *in vivo* mutagenesis could expand the genetic diversity of CelB and improve the system’s evolutionary outcomes (Figure 1B).

By integrating mutagenesis, selection, and enrichment into a single automated process, we achieved efficient and high-throughput evolution of CelB. The continuous culture system supported the evolution of large variant libraries (>10^9^ evolving cells per culture), enabling rapid optimization of CelB with minimal manual intervention.

### Screening and characterization of CelB variants

After completing the evolution experiment, the culture was screened using a blue-white screening method on LB agar plates containing X-gal and aTc. Dark-blue colonies, indicative of higher CelB activity, were selected and grown in LB medium with aTc to induce CelB expression. Following incubation with the inducer, cells were heated at 75^°^C for 15 minutes before performing the chlorophenol red-β-D-galactopyranoside (CPRG) assay in 96-well plates to confirm their beta-galactosidase activity. This heat treatment served two purposes: it selected for thermostable CelB variants and disrupted the bacterial cell envelopes, allowing CPRG penetration for enzymatic activity assays.

Ten variants showing relatively high activity in the CPRG assay are shown in Figure 2 A. Most of them showed higher activity than the wild-type CelB. Compared to wild-type CelB, three variants including AA10, W10 and T1, exhibited 70% increase of enzymatic activity in the heat-treated cells. The mutations of the ten were identified by DNA Sanger sequencing and many of them shared the same mutations (Figure 2B). Three variants, AA10, M10 and T1 share the identical mutations, including G72E and E365K, while M7 only acquired one mutation in E365K. Another two variants, W10 and W20, share the same mutations in L412W, Y416H, and K436N. B5-2 and W21 also shared the same mutation M424T. There were not any mutation found at the promoter region for all variants except W21 that acquired an additional mutation at the *tetO* operon that may transcriptionally increase protein expression.

**Figure 2.**
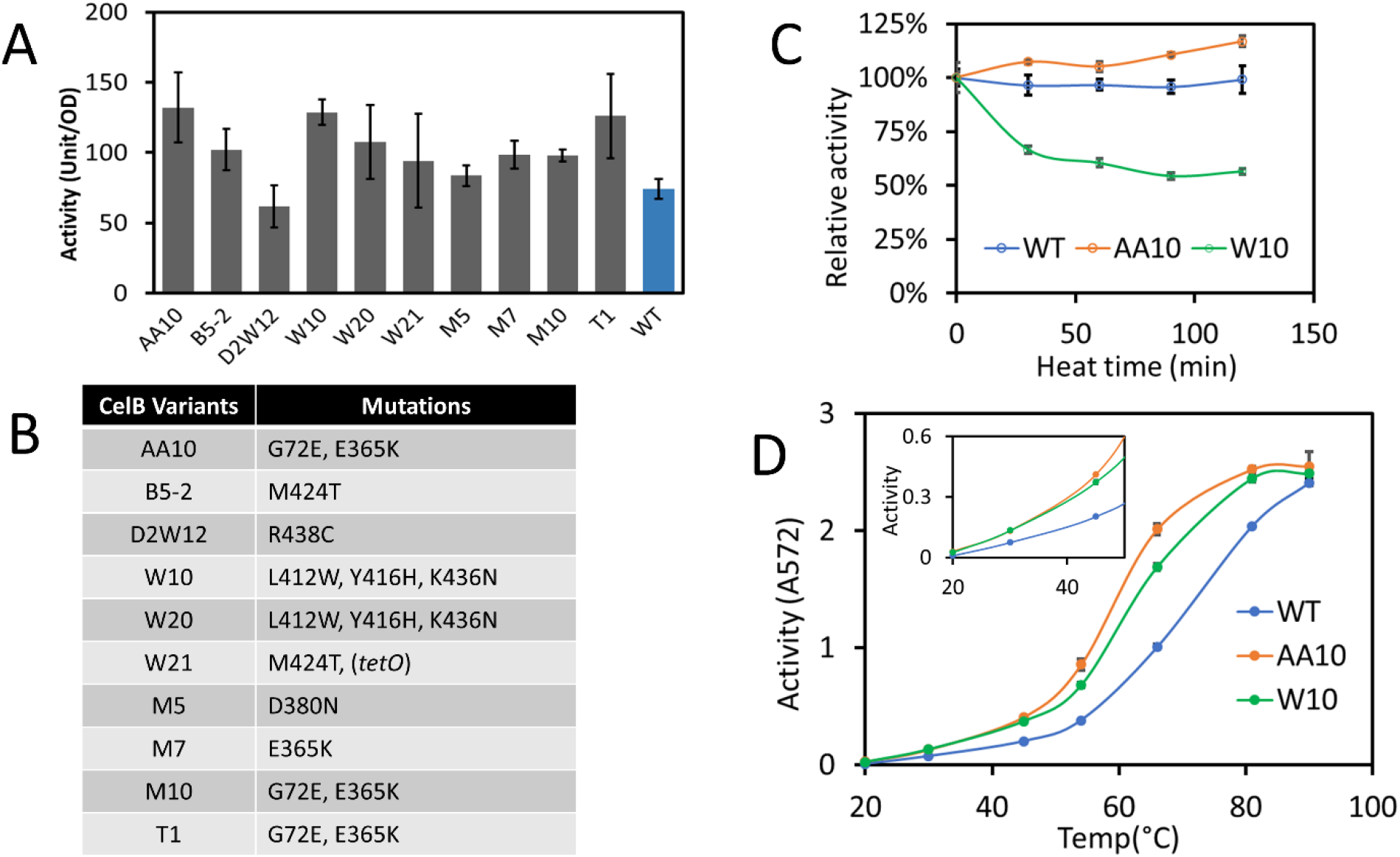
Characterization of CelB variants obtained from continuous directed evolution. (A) CPRG assay results for CelB variants selected from blue-white screening. (B) Amino acid mutations in CelB variants, as identified by DNA sequencing. (C) Thermostability of CelB proteins following heat treatment at 80^°^C over time. Enzymatic activities, represented by the slopes of CPR production curves, were measured at 25^°^C after each heat treatment and subsequently normalized to the initial activity observed without heat exposure. All proteins were overexpressed and purified before testing. (D) Enzymatic activity of wild-type CelB and two variants across a range of temperatures. The inset provides a detailed view of activity levels below 50^°^C. Data points represent means ± standard deviation of four replicates. Data are presented as means ± standard deviation of at least four replicates.

We further characterized the activities of the two top-performing variants, AA10 and W10, under various conditions. To assess their thermostability, the purified proteins were heat-treated at 80^°^C for varying time intervals before testing their enzymatic activity at room temperature (Figure 2C). While the wild-type enzyme maintained consistent activity, in agreement with previous findings (13–15), the two variants behaved differently. AA10 exhibited a slight increase in activity upon heat treatment, while W10 lost approximately 40% of its activity. Additionally, the activity of the variants was evaluated across a range of temperatures (Figure 2D). We observed that, as the temperature increased from 20^°^C to 90^°^C, the activity of the wild type rose rapidly. In contrast, the two variants reached a plateau in activity around 80^°^C, suggesting that our evolution approach has shifted their optimal temperature for activity. Although the two CelB variants showed similar activity to the wild type at 90^°^C, they displayed significantly higher activity at lower temperatures (30^°^C to 80^°^C). Our results demonstrate that the GCCDE approach effectively evolved and selected CelB variants that retained thermostability while exhibiting enhanced enzymatic activity at lower temperatures. Despite the wild-type CelB being considered highly optimized for stability and flexibility (15), our ability to engineer CelB variants with the desired performance in a single evolutionary experiment underscores the robustness and efficiency of this method. These findings highlight the potential of GCCDE to achieve protein engineering outcomes, even for proteins believed to be near their evolutionary limits.

### Kinetic study and protein structure analysis

We next investigated the enzymatic kinetic parameters and mutations of AA10 and W10. The kinetic parameters of these variants were compared to those of the wild-type CelB (Figure 3A). Both AA10 and W10 achieved an approximately 30% increase of *k*_cat_, the turnover number of the enzyme. Additionally, both mutants showed increased binding affinity toward the substrate, as indicated by lower *K*m values. Consequently, the overall catalytic efficiency (*k*_cat_/*K*m) for each variant increased twofold or more compared to the wild type at room temperature.

**Figure 3:**
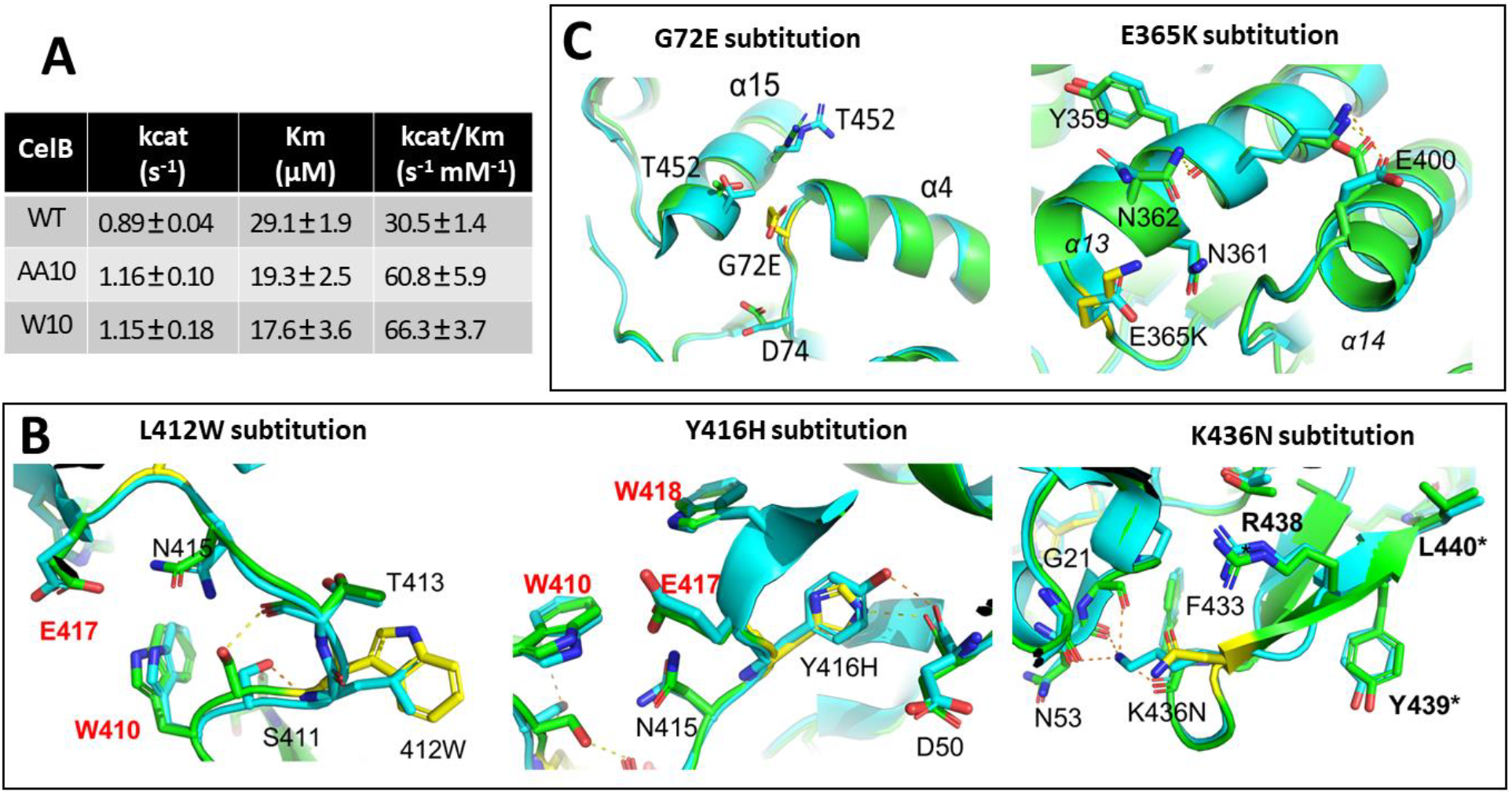
Kinetic parameters and structural analysis of CelB proteins. (A) The kinetic parameters of the wild-type CelB and two variants (AA10 and W10), measured at 25^°^C using CPRG as substrate. (B) Structural alignment of the wild type and W10. (C) Structural alignment of the wild type and AA10. Note: The wild-type CelB (PDB id: 3APG) is in cyan while variants are in green. The structures of variants were predicted from AlphaFold2. Residues in yellow color indicate amino acid substitutions. Active site residues are labeled in bolded red text, and inter-subunit residues are indicated by bolded text with an asterisk. Yellow and orange dashed lines indicate the hydrogen bonds formed between residues in variants and wild type, respectively. The secondary structure element, α-helices (α) are indicated in gray italics.

Amino acid substitutions in the two CelB variants are illustrated in the multiple sequence alignment with the wild-type protein (Figure S1). The W10 variant harbors three key substitutions: L412W, Y416H, and K436N (Figure 3B) that are located at the C-terminal region, which contains several well-conserved active-site residues and subunit-interface residues (15). To better explain the role of the mutations, the three-dimensional structure of W10 was predicted by AlphaFold2 (16) and compared to the structure of the wild type protein that is publicly available (PDB id: 3APG). The substitution of L412 with a bulkier tryptophan near the active residue W410 likely facilitates substrate binding by reducing the size of the binding pocket. Additionally, the Y416H substitution could induce spatial shifts in the adjacent active residues including E417 and W418 (15), potentially promoting both galactose binding and catalytic efficiency of the enzyme (17). Lastly, K436N, located on the protein’s exterior near inter-subunit contact regions, may influence the protein’s overall thermostability. Replacing the charged lysine with an uncharged asparagine is predicted to disrupt the hydrogen bonds with G21, N53 and F433. This mutation also switch Y439 and Y438, the residues for interfacial interactions and the homotetramer formation of CelB (15). Consequently, the K436N substitution may impair enzyme rigidity, which could explain the decreased stability of W10 variant under heat treatment (Figure 2C).

The AA10 variant contains two substitutions: G72E and E365K. Although the two mutations does not occur within the catalytic active sites, the resulting alternations in secondary structures could influence inter-subunit interactions that are critical for substrate entry and/or product release (18). These changes likely contribute to the enhanced catalytic activity and substrate binding affinity observed in the kinetic study (Figure 3A). The glycine residue at position 72 is highly conserved across species (15), and replacing it with the charged glutamic acid introduces electrostatic attraction with R448 on the α15 helix (Figure 3C). Additionally, the larger glutamic acid residue may introduce steric clashes to the local pocket, altering the arrangement of nearby residues, including T452 and D74. Consequently, the conformation of the α15 helix and the adjacent loop undergo significant rearrangements (Figure 3B). The second substitution, E365K, reverses the charge at its position, which may induce conformational changes in the α13 and α14 helices. Collectively, these structural shifts in helices, including α13, α14, and α15, along with potential changes in neighboring helices, loops, and beta-sheets, could significantly impact the overall architecture of the enzyme. These structural changes could enhance inter-subunit interactions, stabilize the quaternary structure, and promote the formation of homotetramers. Such rearrangements likely improve the cooperative dynamics and catalytic efficiency of CelB, ultimately enhancing its enzymatic performance.

## CONCLUSION

We have developed a growth-coupled continuous protein engineering approach that automates mutagenesis, selection, and enrichment simultaneously. Using this method, we successfully engineered the thermostable enzyme CelB to exhibit enhanced β-galactosidase activity at lower temperatures while maintaining its thermal stability, all within a single evolutionary experiment. This success demonstrates the effectiveness of our growth-coupled continuous directed evolution (GCCDE) approach. GCCDE enables genetic diversification, selection, and enrichment in a single process, significantly boosting the efficiency of directed evolution. By integrating *in vitro* and *in vivo* mutagenesis, our method expands genetic diversity and improve evolutionary outcomes. Coupling enzymatic activity to cellular growth enables high-throughput selection of improved variants. With a continuous culture system and the MutaT7 platform, we achieved automated enzyme engineering with high-throughput and real-time selection exceeding 10^9^ cells in only one bioreactor. The resulting CelB variants showed higher β-galactosidase activity at room temperature while retaining their thermal stability. Key mutations were identified that likely improved substrate binding at lower temperatures while preserving overall structural integrity. This GCCDE approach is broadly applicable for engineering any protein whose activity can be linked to cellular growth, offering a powerful solution for high-throughput protein optimization and evolution.

## MATERIALs AND METHODS

### Strains and growth conditions

*E. coli* NEB 10-beta cells (NEB, cat# C3019H) were used for plasmid construction, and *E. coli* JM109 cells were utilized for protein overexpression in Lysogeny Broth (LB) medium. NEB SOC outgrowth medium (NEB, cat# B9020S) was used for recovery following transformations. The *E. coli* Dual7 strain (10) , derived from DH10B and lacking native β-galactosidase activity, was used for *in vivo* directed evolution experiments.

Lactose minimal medium, modified from M63 medium, contained lactose (10 g/L), (NH_4_)_2_SO_4_ (2 g/L), KH_2_PO_4_ (20 g/L), K_2_HPO_4_ (48 g/L), FeSO_4_ (0.5 mg/L), thiamine (0.5 mg/L), MgSO_4_ (1 mM), NaCl (0.1 g/L), tryptone (0.2 g/L), and yeast extract (0.1 g/L), with pH adjusted to 7.0–7.2. M63-glucose medium was similarly prepared but substituted lactose with glucose (2 g/L). Kanamycin (50 µg/ml) was added when required.

### Plasmid construction and CelB library preparation

Plasmids were constructed using the NEBuilder® HiFi DNA Assembly kit (NEB), and PCR reactions were performed with NEB Q5 High-Fidelity Polymerase. The *celB* gene from *Pyrococcus furiosus* (14) was synthesized by IDT and cloned into the pQE80 plasmid. The T5 promoter in pQE80 was modified by replacing one lacO sequence with tetO (5’-tccctatcagtgatagaga-3’) and mutating the second lacO to enable regulation solely by TetR and anhydrotetracycline (aTc). The CelB gene, engineered hybrid promoter, and TetR gene from pDA303 (19) were assembled to create plasmid pDA381. The low-copy-number pDA386 was then derived from pDA381 by replacing its origin of replication with pSC101 Ori from pLC-F (20). Plasmid pDA386 served as the vehicle for *in vivo* mutagenesis and as the template for error-prone PCR. All the primers used in this study were in Table S1 in the supplementary material.

Error-prone PCR was performed to generate a starting library targeting CelB’s catalytic region (amino acids 350-472) (13, 15, 17). The reaction mixture (40 µl) contained 0.6 ng/µl template DNA, 0.2 mM dATP/dGTP, 1 mM dCTP/dTTP, 5 mM MgCl_2_, 0.2 mM MnCl_2_, 0.05 U/ml NEB Taq polymerase, and 0.4 µM primers (386-mut-F349 and 381-V1-R). PCR conditions were: initial denaturation at 94^°^C for 2 min; 18 cycles of 94^°^C for 20 s, 50^°^C for 30 s, and 68^°^C for 30 s; followed by a final extension at 68^°^C for 5 min. The resulting PCR product was verified by agarose gel electrophoresis, purified, and assembled into plasmids. These plasmids were transformed into *E. coli* Dual7 cells (10) using a previously established method (22), creating the CelB starting library. The library was recovered in 2 ml of SOC medium before inoculating to 15 ml of lactose minimal medium with kanamycin and aTc (20ng/ml) for *in vivo* continuous evolution.

### Growth-coupling continuous directed evolution and selection

The starting CelB library culture (17 ml) was grown at 37°C in the Chi.Bio continuous culture system (23). When the OD_600_ reached 0.8, the turbidostat was activated to maintain this level overnight by supplying fresh medium as needed. Subsequently, the OD_600_ was controlled at 0.6 (∼10^8^ CFU/ml, mid-exponential phase). The evolution experiment continued for two weeks, with approximately 1.7 × 10^9^ cells undergoing *in vivo* mutagenesis in the bioreactor.

Lactose in the medium activated MutaT7 expression, driving continuous mutagenesis of the *celB* gene downstream of the T7 promoter on the plasmids. Variants with higher enzymatic activity hydrolyzed lactose more efficiently, promoting faster bacterial growth. Slower-growing cells were gradually washed out of the bioreactor. The temperature was gradually reduced from 37^°^C to 27^°^C, selecting for CelB variants with enhanced β-galactosidase activity at lower temperatures. At the end of the evolution experiment, the culture was grown in M63-glucose medium to stop MutaT7 expression and stabilize mutations.

### Screening and measurement of beta-galactosidase activity

The evolved cells were diluted and plated on LB agar plates supplemented with 5-bromo-4-chloro-3-indolyl-beta-D-galactopyranoside (X-gal, 40 µg/ml), aTc (30 ng/ml), and kanamycin (50 µg/ml) for blue-white screening. Dark blue colonies, indicative of high β-galactosidase activity, were selected and grow in LB medium to overexpress CelB with aTc induction. The cells were washed and resuspended in phosphate-buffered saline buffer (PBS, pH 7.4). The suspensions were heated at 75^°^C for 15 minutes to select heat-resistant variants and to lyse the cells, allowing CPRG to enter cells for enzymatic activity measurement.

CelB activity was quantified using the CPRG assay. In this assay, CelB hydrolyzes CPRG to release chlorophenol red (CPR), which was monitored at 572 nm (A_572_) using a microplate reader (Molecular Devices Inc.). CPRG was added to a final concentration of 0.5 mM, and glucose was added to 2.7% to screen for beta-galactosidase variants tolerant to glucose inhibition. Enzymatic activity was calculated based on the slope of the CPR production curve, defined as an increase of 0.001 absorbance units per minute at A_572_. To compare enzymatic activity among variants, the density (OD_600_) of all cultures was measured to normalize enzymatic activity.

### Protein Purification and Characterization

The selected CelB variants were cloned into a high-copy-number plasmid with a 6xHis tag using pDA381 as the backbone. Proteins were overexpressed in E. coli JM109 (DE3) upon aTc induction (30 ng/ml) and purified using Ni-NTA agarose resin (ThermoFisher Scientific, cat# 88221), followed the protocol described previously (19). Purity and molecular weight of the purified proteins were assessed by reducing SDS-PAGE, and protein concentration was determined using a BCA assay (ThermoFisher Scientific, cat# 23252). Enzymatic activity was measured using the CPRG assay described above. Kinetic parameters, including *Km* and *k*_*cat*_, were determined using the classical Eadie-Hofstee linearization plot (24). For thermostability testing, proteins were incubated at 80^°^C in a thermocycler (Bio-Rad Inc.) for various time intervals before conducting the CPRG assay at 25^°^C. Additionally, temperature-dependent activity was evaluated by incubating the purified enzymes with CPRG across varying temperatures for 15 min, followed by absorbance measurements at 572nm (A_572_). Blank controls were prepared by incubating the CPRG solution without enzyme under the same conditions.

### Sequence alignment and structure analysis

Mutations in the selected high-activity variants were identified by DNA sequencing. The amino acid sequences of CelB variants were aligned with the wild-type CelB sequence using Clustal Omega (25) and rendered in EsPript 3 (26). Three-dimensional structural models of the CelB mutants were generated in ColabFold platform (27) with AlphaFold2 (16). Structural visualization and alignment were conducted using PyMOL (Version 2.5.4, Schrödinger, LLC).

## ACKNOWLEDGMENTS

This work was supported by funding from Dartmouth College.

## DATA AVAILABILITY

All data generated during this study are included in this published article and in its supplemental material files. Raw data were available upon request.

